# Integrated multi-omics approach reveals the role of SPEG in skeletal muscle biology including its relationship with myospryn complex

**DOI:** 10.1101/2023.04.24.538136

**Authors:** Qifei Li, Jasmine Lin, Shiyu Luo, Klaus Schmitz-Abe, Rohan Agrawal, Melissa Meng, Behzad Moghadaszadeh, Alan H. Beggs, Xiaoli Liu, Mark A. Perrella, Pankaj B. Agrawal

**Affiliations:** Division of Newborn Medicine, Boston Children’s Hospital, Harvard Medical School; Boston, Massachusetts 02115, USA; Division of Genetics and Genomics, Boston Children’s Hospital, Harvard Medical School; Boston, Massachusetts 02115, USA; The Manton Center for Orphan Disease Research, Boston Children’s Hospital, Harvard Medical School; Boston, Massachusetts 02115, USA; Division of Neonatology, Department of Pediatrics, University of Miami Miller School of Medicine and Holtz Children’s Hospital, Jackson Health System, Miami, FL, USA; Division of Pulmonary and Critical Care Medicine, Brigham and Women’s Hospital, Harvard Medical School; Boston, Massachusetts 02115, USA; Department of Pediatric Newborn Medicine, Brigham and Women’s Hospital, Harvard Medical School; Boston, Massachusetts 02115, USA

## Abstract

Autosomal-recessive mutations in *SPEG* (striated muscle preferentially expressed protein kinase) have been linked to centronuclear myopathy. Loss of SPEG is associated with defective triad formation, abnormal excitation-contraction coupling, and calcium mishandling in skeletal muscles. To elucidate the underlying molecular pathways, we have utilized multi-omics tools and analysis to obtain a comprehensive view of the complex biological processes. We identified that SPEG interacts with myospryn complex proteins (CMYA5, FSD2, RyR1), and SPEG deficiency results in myospryn complex abnormalities. In addition, transcriptional and protein profiles of SPEG-deficient muscle revealed defective mitochondrial function including aberrant accumulation of enlarged mitochondria on electron microscopy. Furthermore, SPEG regulates RyR1 phosphorylation at S2902, and its loss affects JPH2 phosphorylation at multiple sites. On analyzing the transcriptome, the most dysregulated pathways affected by SPEG deficiency included extracellular matrix-receptor interaction and peroxisome proliferator-activated receptors signaling, which may be due to defective triad and mitochondrial abnormalities. In summary, we have elucidated the critical role of SPEG in triad as it works closely with myospryn complex, phosphorylates JPH2 and RyR1, and demonstrated that its deficiency is associated with mitochondrial abnormalities. This study emphasizes the importance of using multi-omics techniques to comprehensively analyze the molecular anomalies of rare diseases.

**Synopsis:** 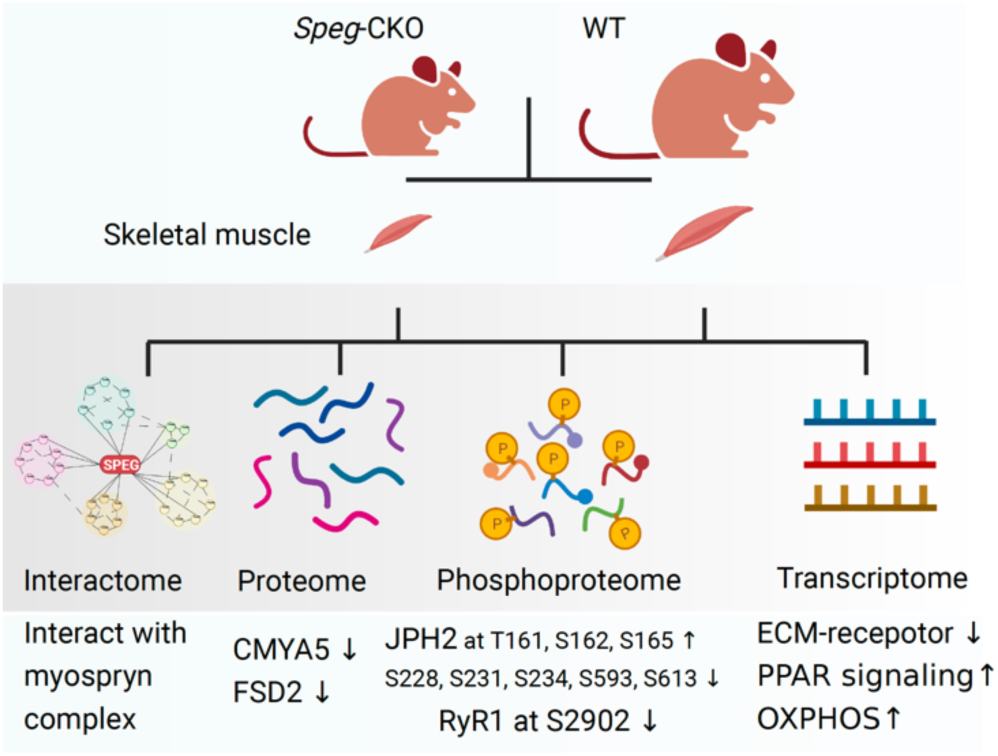

We have previously linked mutations in *SPEG* (striated preferentially expressed protein) with a recessive form of centronuclear myopathy and/or dilated cardiomyopathy and have characterized a striated muscle-specific SPEG-deficient mouse model that recapitulates human disease with disruption of the triad structure and calcium homeostasis in skeletal muscles. In this study, we applied multi-omics approaches (interactomic, proteomic, phosphoproteomic, and transcriptomic analyses) in the skeletal muscles of SPEG-deficient mice to assess the underlying pathways associated with the pathological and molecular abnormalities.

- SPEG interacts with myospryn complex proteins (CMYA5, FSD2, RyR1), and its deficiency results in myospryn complex abnormalities.
- SPEG regulates RyR1 phosphorylation at S2902, and its loss affects JPH2 phosphorylation at multiple sites.
- SPEGα and SPEGβ have different interacting partners suggestive of differential function.
- Transcriptome analysis indicates dysregulated pathways of ECM-receptor interaction and peroxisome proliferator-activated receptor signaling.
- Mitochondrial defects on the transcriptome, proteome, and electron microscopy, may be a consequence of defective calcium signaling.

## Introduction

Congenital myopathies (CM) present with skeletal muscle weakness and hypotonia of varying severity, time of onset, genetic cause, and often subclassified by pathological findings. Centronuclear myopathy (CNM) is a common subtype characterized by the mislocalization of nuclei to the center of myofiber instead of the periphery. The clinical spectrum of CNM is diverse among affected individuals, and about 60%-80% of CNM can be explained by mutations in *MTM1* (myotubularin 1) (1), *DNM2* (dynamin 2) (2), *BIN1* (bridging integrator 1) (3), *RyR1* (ryanodine receptor 1) (4), *CACNA1S* (Voltage-dependent L-type calcium channel subunit alpha-1S) (5), and *SPEG* (striated preferentially expressed gene) (6).

We have previously linked recessive variants in *SPEG* with CNM, CM or dilated cardiomyopathy (DCM), CNM and DCM often presenting together (6, 7). SPEG is a member of the myosin light chain kinase (MLCK) protein family, important for myocyte function and the regulation of the actin-based cytoskeleton (8). We have determined that SPEG localizes in a double line, in alignment with the terminal cisternae of the sarcoplasmic reticulum (SR) of the triad (9). Skeletal excitation-contraction (E-C) coupling and calcium homeostasis require triads, the nanoscopic microdomains formed adjacent to Z-lines by apposition of transverse tubules and terminal cisternae of the SR (10). Furthermore, using striated muscle-specific *Speg* knockout (*Speg*-CKO) mice, we have shown that SPEG deficiency is associated with defective triad formation leading to abnormal E-C coupling, and calcium mishandling in skeletal muscles (9). Triad abnormalities are also seen with DNM2 (11), MTM1 (12) and BIN1 (13) proteins that are associated with CNM, and we have previously demonstrated that SPEG interacts with MTM1 (6) and DNM2 (14). The molecular relationship of SPEG with triadic proteins and the effects of SPEG deficiency on downstream pathways need to be elucidated.

Here, we utilized mouse model of SPEG deficiency and applied multi-omics approaches (interactomic, proteomic, phosphoproteomic, and transcriptomic analyses) to determine interacting proteins and molecular pathways associated with SPEG. We identified members of the myospryn complex that include cardiomyopathy-associated protein 5 (CMYA5 or myospryn), fibronectin type III and SPRY domain-containing protein 2 (FSD2 or minispryn), and ryanodine receptor 1 (RyR1)) as novel SPEG-interacting partners using mass spectrometry-based protein–protein interaction analysis, and further confirmed by co-immunoprecipitation (co-IP) assays. In addition, SPEG phosphorylates multiple sites of JPH2 and RyR1 (S2902 in particular), and its deficiency causes marked reduction in CMYA5, FSD2, and MTM1 protein levels. Furthermore, transcriptomic analysis indicates dysregulated pathways of extracellular matrix (ECM)-receptor interaction suggestive of defective focal adhesion. Mitochondrial dysfunction due to SPEG deficiency was noted by transcriptomic analysis and EM.

## Results

### Levels of SPEGα and SPEGβ increase over time

SPEG locus encodes for four distinct tissue-specific isoforms: SPEGα (260 kDa) and SPEGβ (350 kDa) are highly expressed in skeletal and cardiac muscles, while APEG1 in vascular tissues, and BPEG in the brain and aorta (8). To evaluate the relative expression of SPEGα and SPEGβ over different time points, we performed quantitative real time PCR and immunoblot in mouse quadriceps muscles and found that the transcript and protein levels of SPEGα and SPEGβ gradually increased from embryonic day 18.5 to 2 months of age (**Figure 1A and 1B**). To assess the role of SPEG in adult skeletal muscle, 2-month-old WT and *Speg*-CKO mice were used for the following experiments.

**Figure 1.**
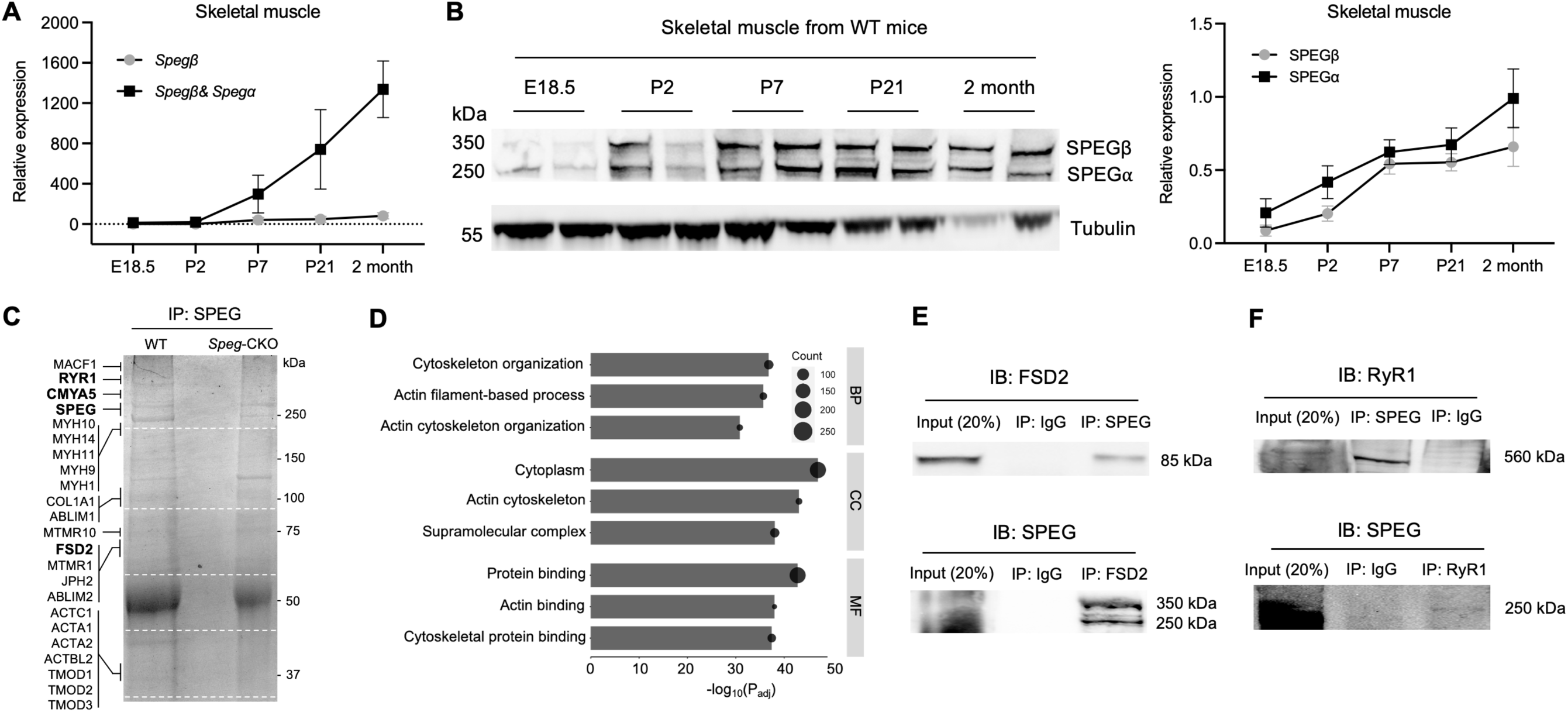
SPEG interacts with myospryn complex proteins in the skeletal muscles. (A) Quantitative real-time PCR analysis of *Speg α* and *β* mRNA expression in the skeletal muscles of wild-type mice. *Speg α and β mRNA expression at* P2 was used for normalization. (B) Immunoblot analysis for SPEGα and SPEGβ isoforms and quantification. Tubulin is used as a loading control. E: embryonic; P: postnatal. Experiments were performed using at least three different biological samples. (C) Identification of SPEG-binding partners in skeletal muscle using Coomassie blue-stained gel after elution of SPEG–immune complexes. White dashed lines indicate gel fragments used for mass spectrometry analysis. IP, immunoprecipitation. (D) Top three gene ontology enrichments of BP, CC, and MF using the SPEG-binding partners in skeletal muscle. MF: molecular function; BP: biological process; CC: cellular component. (E) SPEGβ and SPEGα and FSD2 (fibronectin type III and SPRY domain-containing protein 2) coimmunoprecipitated from skeletal muscle lysates with the use of rabbit anti-SPEG generated against a FLAG-tagged aortic preferentially expressed gene-1 fusion protein and anti-FSD2 antibodies. (F) SPEGα and RyR1 coimmunoprecipitated from skeletal muscle lysates with the use of rabbit anti-SPEG and anti-RyR1 antibodies.

### SPEG interacts with myospryn complex proteins in skeletal muscle

Pooled skeletal muscles (quadriceps, gastrocnemius, and triceps) from WT and *Speg*-CKO mice were used to determine the binding partners of SPEG using immunoprecipitation (**Figure 1C**) followed by mass spectrometry assay. A total of 340 proteins were found to be potential SPEG-binding partners after comparing the abundance of protein fragments between WT and *Speg*-CKO. Among these 340 proteins, 193 of them were fully absent from *Speg*-CKO, and the resting 147 proteins in WT are more than 1.6-fold higher than those in *Speg*-CKO. These binding partners were then analyzed by Gene Ontology enrichment, and the top three GO enrichments of biological process (BP), cellular component (CC), and molecular function (MF) were listed in **Figure 1D**. Interestingly, cytoskeleton organization and protein binding associated with actin filament were highly enriched among these potential binding partners. SPEG has been reported to interact with MTM1 (6), desmin (15), and DNM2 (14) in skeletal muscle. In these binding proteins MTM1 and DNM2 were not found, but MTMR1, MTMR10, desmin, and JPH2 were detected. Additionally, we found all three proteins (CMYA5, FSD2, RyR1) that formed the myospryn complex in the terminal SR of triads which has been shown abundant in the mass spectrometry findings (16). Using co-IP assays, we found that while both SPEGβ (350 kDa) and SPEGα (250 kDa) co-IP with FSD2 (**Figure 1E**), whereas only SPEGα co-IP with RyR1 (**Figure 1F**). These findings suggest SPEG proteins are novel binding partners of myospryn complexes in skeletal muscle.

### Loss of SPEG causes myospryn complex and MTM1 abnormalities in skeletal muscles

SPEG engages in a functional network that is critical to triad development and function by regulating the activities of triadic proteins (9). To assess proteomic changes associated with its absence, quadriceps muscles from adult *Speg*-CKO (n =3) and WT mice (n =4) were lysed and processed. Principal component analysis (PCA) in **Figure 2A** showed that *Speg*-CKO groups were distinct from the WT, and volcano plots (**Figure 2B**) display the differentially expressed proteins (DEPs; |FC| > 1.5, *P < 0.05). Noticeably, proteins associated with myospryn complex (CMYA5 and FSD2) were significantly reduced among these 38 dysregulated DEPs (**Figure 2C**), which were enriched in cellular components of cytoplasm, mitochondria, peroxisome, and sarcomere by GO analysis (**Figure 2D**). To further validate the findings, immunoblot and quantification of selected DEPs from the skeletal muscles of WT, *Speg*-CKO and *Mtm1*-KO mice were performed and compared (**Figure 2E**). Immunoblot findings confirmed that loss of SPEG caused significant reduction of CMYA5, FSD2, and MTM1 levels in skeletal muscles. In contrast, the protein levels of CMYA5, FSD2, and SPEG were comparable between *Mtm1*-KO and control mice, suggesting that this reduction in myospryn complex proteins is unique to SPEG deficiency.

**Figure 2.**
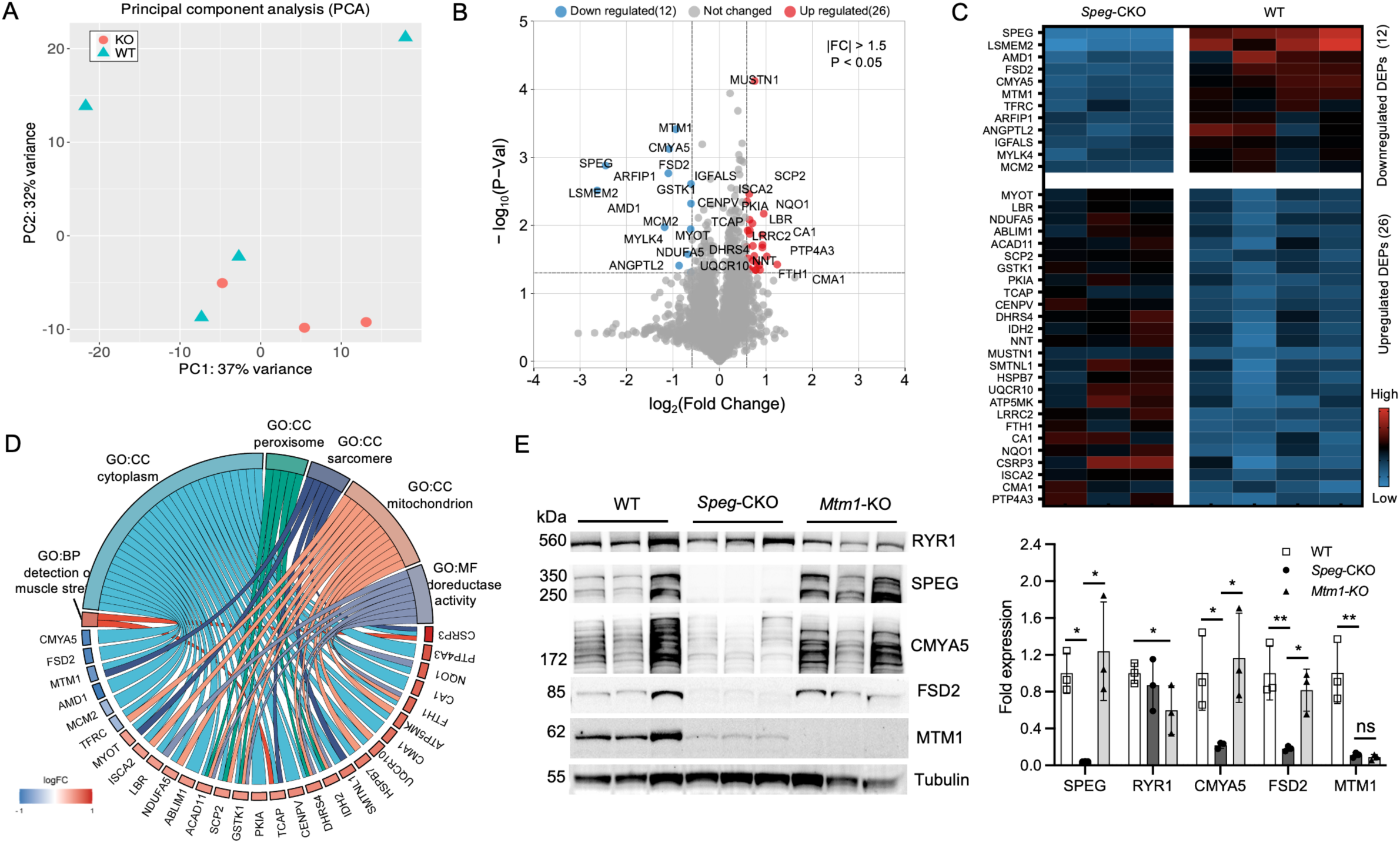
Proteome profiling in the skeletal muscle of *Speg*-CKO mice. (A) Principal component analysis of proteome data. The first and second axes are represented. Colored symbols represent genotypes for each mouse. (B) Volcano plots representing the differentially expressed proteins (DEPs). Upregulated proteins are in red, and downregulated proteins are in blue (P <0.05 and a fold change beyond ±1.5). (C) Heat map of the proteins that were differentially expressed in the skeletal muscle of *Speg*-CKO mice. (D) Chord plot displaying the enrichment analyses of DEPs for GO. (E) Immunoblot analysis and quantification of SPEG, RyR1, CMYA5, FSD2, and MTM1 relative to the expression of tubulin in skeletal muscles of WT, *Speg*-CKO, and *Mtm1*-KO mice (ns, not statistically significant, *P ˂ 0.05, **P ˂ 0.01, n = 3 per genotype; 1-way ANOVA with Tukey’s post hoc test).

### Protein expression profiles reveal mitochondrial abnormalities in SPEG-deficient mice

To further delineate the effects of SPEG deficiency in skeletal muscles, we performed unbiased gene set enrichment analysis (GSEA) using the proteome data from WT and *Speg*-CKO adult quadriceps. A total of 3517 protein sets categorized by gene ontology were visualized using the enrichment map visualization method (17). The red and blue nodes indicate the upregulated and downregulated protein sets, respectively. In SPEG-deficient mice, mitochondria-related and contractile fiber protein sets were significantly upregulated, whereas nucleic acid metabolism and ECM structural protein sets were downregulated (**Figure 3A**). Select GSEA plots displaying the downregulated and upregulated protein sets in *Speg*-CKO mice are shown in **Figure 3B**. The bar charts in **Figure 3C** show the top 5 normalized enrichment score (NES) of upregulated protein sets in *Speg*-CKO proteome, most of which are associated with mitochondria function.

**Figure 3.**
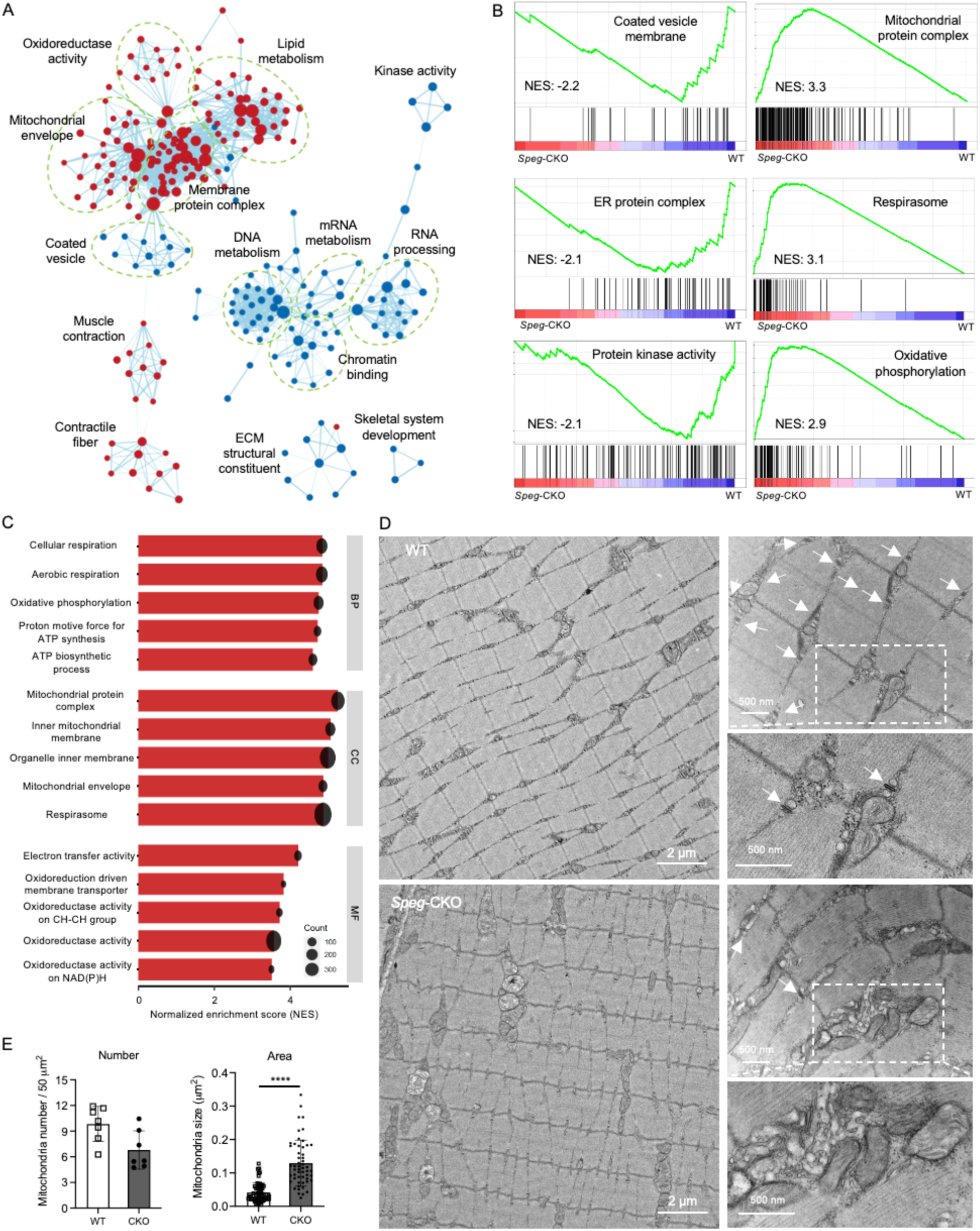
Protein expression profiles in the skeletal muscle of *Speg*-CKO mice. (A) The enrichment map summarizing the result of GSEA of the *Speg*-CKO skeletal muscle proteome, presenting the downregulated (blue dots) and upregulate (red dots) protein sets in *Speg*-CKO mice, comparing with WT mice. The size of a node indicates the size of each protein set, and the edge means two connected nodes share same proteins. (B) The selected GSEA plots presenting the downregulated and upregulate protein sets in *Speg*-CKO mice. (C) The bar charts showing the top 5 NES of upregulate protein sets in *Speg*-CKO proteome. (D) Electron micrographs in quadriceps specimens obtained from WT and *Speg*-CKO mice. The left panel shows an overall organization of muscle structure, and the right panel shows an enlarged view of triad (white arrows) and mitochondria (box) ultrastructure from each group. (E) Quantification of mitochondria number (left bar graph) and area (right bar graph) (****P < 0.0001, n = 3 per group; unpaired 2-tailed t test).

### SPEG-deficient muscle exhibits accumulated swollen mitochondria

To further assess the effects of SPEG deficiency on mitochondria morphology, sections of quadriceps muscle were examined using electron microscopy (EM). In **Figure 3D**, the *Speg*-CKO muscle displayed structural triad anomalies (arrows) with regions of disoriented or absent triads, twisted Z-lines, and abnormally accumulated swollen mitochondria (square box). Quantitative analysis of EM findings between *Speg*-CKO and WT muscles revealed that there was a statistically significant increase (****P < 0.0001) in the mitochondrial area of SPEG-deficient mice while the number of mitochondria remained relatively unchanged (**Figure 3E**). Additionally, immunoblot analysis of mitochondrial dynamics and OxPhos proteins was performed, and there was no difference between *Speg*-CKO and WT muscles (**Supplemental Figure 1**). Overall, these results indicate that SPEG depletion results in aberrant mitochondrial morphology in skeletal muscle.

### SPEG phosphorylates multiple phosphorylation sites on JPH2 in skeletal muscle

The SPEGα and SPEGβ proteins contain variable Ig-like (SPEGβ: 9 Ig-like; SPEGα: 7 Ig-like), three fibronectin type III, and two tandemly arranged serine/threonine kinase domains (8). It has been reported that in cardiac muscle only SPEGα binds with JPH2 (18), which is expressed in both skeletal and cardiac muscles and stabilizes cardiac dyads between T-tubule and junctional SR membranes, ensuring appropriate intracellular calcium signaling (19, 20). Additionally, the first kinase of SPEG can phosphorylate JPH2, while its phosphorylation sites are still to be determined (21). Nevertheless, the kinase function of SPEG in skeletal muscle is unknown. Here, we applied a phosphoproteomics approach to determine the phosphorylation substrates of SPEG in skeletal muscle. The PCA in **Figure 4A** displays that *Speg*-CKO versus the WT groups, and volcano plots (**Figure 4B**) illustrates the differentially expressed phosphosites (DEpPs). A total of 282 dysregulated DEpPs (129 upregulated and 153 downregulated; |FC| > 1.5, *P < 0.05) were detected as shown in **Figure 4C**. These DEpPs could be directly or indirectly regulated by SPEG deficiency. To identify potential DEpPs directly regulated by SPEG, 22 overlapping proteins were selected (**Figure 4D**) based on our identified SPEG-binding partners (**Figure 1C**). Among these 22 proteins, the protein levels of SPEG and CMYA5 have been found to be significantly decreased (**Figure 2E**), which likely result in their reducing levels of phosphorylation. For the rest of protein levels, either they are no significant change or |FC| < 1.5 (SPTB). **Figure 4E** displays a heat map of selected DEpPs that were statistically significant (*P < 0.05; **P < 0.01) in relation to their corresponding protein level. It is noteworthy that multiple DEpPs (T161, S162, S165, S228, S231, S234, S593, and S613) of JPH2 were up- and down-regulated (**Figure 4F**). JPH2 is a known binding protein and phosphorylation substrate of SPEG in cardiac muscles (18), and here we identified multiple phosphorylation sites of JPH2 were deregulated after SPEG deficiency in skeletal muscle. Additionally, other phosphorylation sites of junctional SR proteins (RyR1-pS2902, CACNA2D1-pS119, MTMR10-pS603, and PHKA1-(pS973; pS982)) and filament proteins (MYH1-(pS36; pY413), MYBPC2-pS476, TNNI2-pS118, and XIRP2-pS2390) were also affected by loss of SPEG.

**Figure 4.**
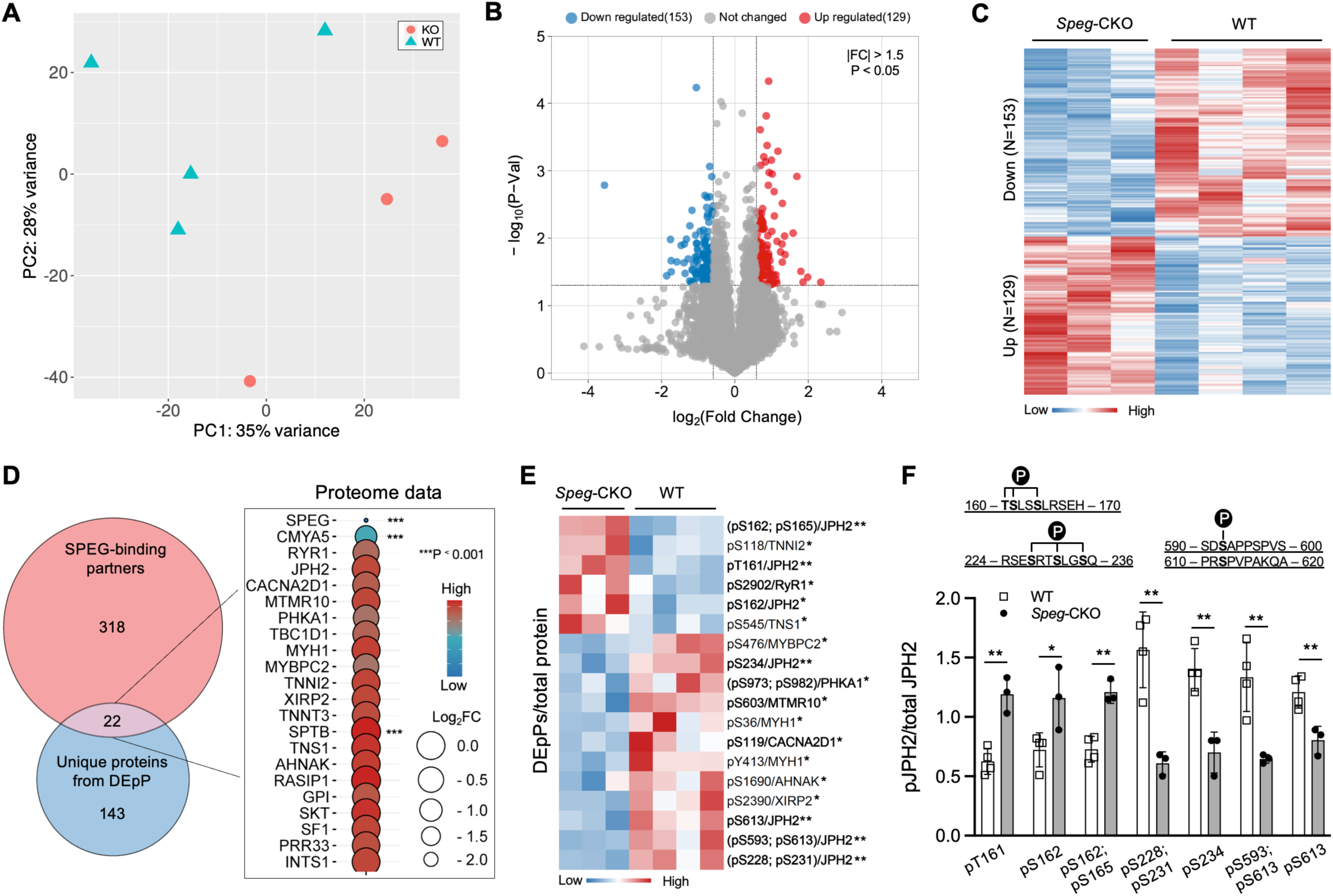
Phosphoproteome profiling in the skeletal muscle of *Speg*-CKO mice. (A) Principal component analysis of phosphoproteome data. The first and second axes are represented. Colored symbols represent genotypes for each mouse. (B) Volcano plots representing the differentially expressed phosphosites (DEpPs). Upregulated DEpPs are in red, and downregulated DEpPs are in blue (P <0.05 and a fold change beyond ±1.5). (C) Heat map of the DEpPs in the skeletal muscle of *Speg*-CKO mice. (D) Left panel: Venn plot of SPEG-binding partners identified in Figure 1C and unique proteins from DEpPs. Right panel: overlapping protein expression identified from proteome analysis. (E) Heat map of selected DEpPs relative to their corresponding protein level. Bold DEpPs: associated with jSR protein and function; other DEpPs: associated with filaments. (F) DEpP quantification of JPH2 at T161, S162, S165, S228, S231, S234, S593, and S613 normalized to total JPH2 protein level (*P < 0.05; **P < 0.01; unpaired 2-tailed t test).

### SPEG phosphorylates S2902 on RyR1 in skeletal muscle

The SPEG kinase has been reported to phosphorylate both RyR2^S2367^ (22) and SERCA2a^T484^ (21) in cardiac muscle, which inhibits diastolic calcium release and promote calcium reuptake into the SR, regulating calcium homeostasis between cytosol and SR. RyR1 is part of the myospryn complex and it was noted to be reduced but the reduction was not statistically significant (**Figure 2E**). The phosphorylation levels of RyR1 were evaluated and several RyR1 phosphorylation sites were detected, listed in **Figure 5A**. Interestingly, one of those, RyR1 S2902 indicated a statistically significant difference (pRyR1^S2902^/total RyR1; *P < 0.05) confirmed by quantification analysis (**Figure 5B**). To further verify this finding, a custom-made phospho-epitope specific polyclonal antibody that recognizes RyR1 phosphorylated S2902 was generated and immunoblot analysis was performed (**Figure 5C**). The RyR1 phosphorylation at S2902 was considerably decreased (*P < 0.05) in *Speg*-CKO mice after correction for total RyR1 level between *Speg*-CKO and WT mice (**Figure 5D and 5E**). Thus, these data confirm that SPEG depletion leads to decreased RyR1 phosphorylation at S2902.

**Figure 5.**
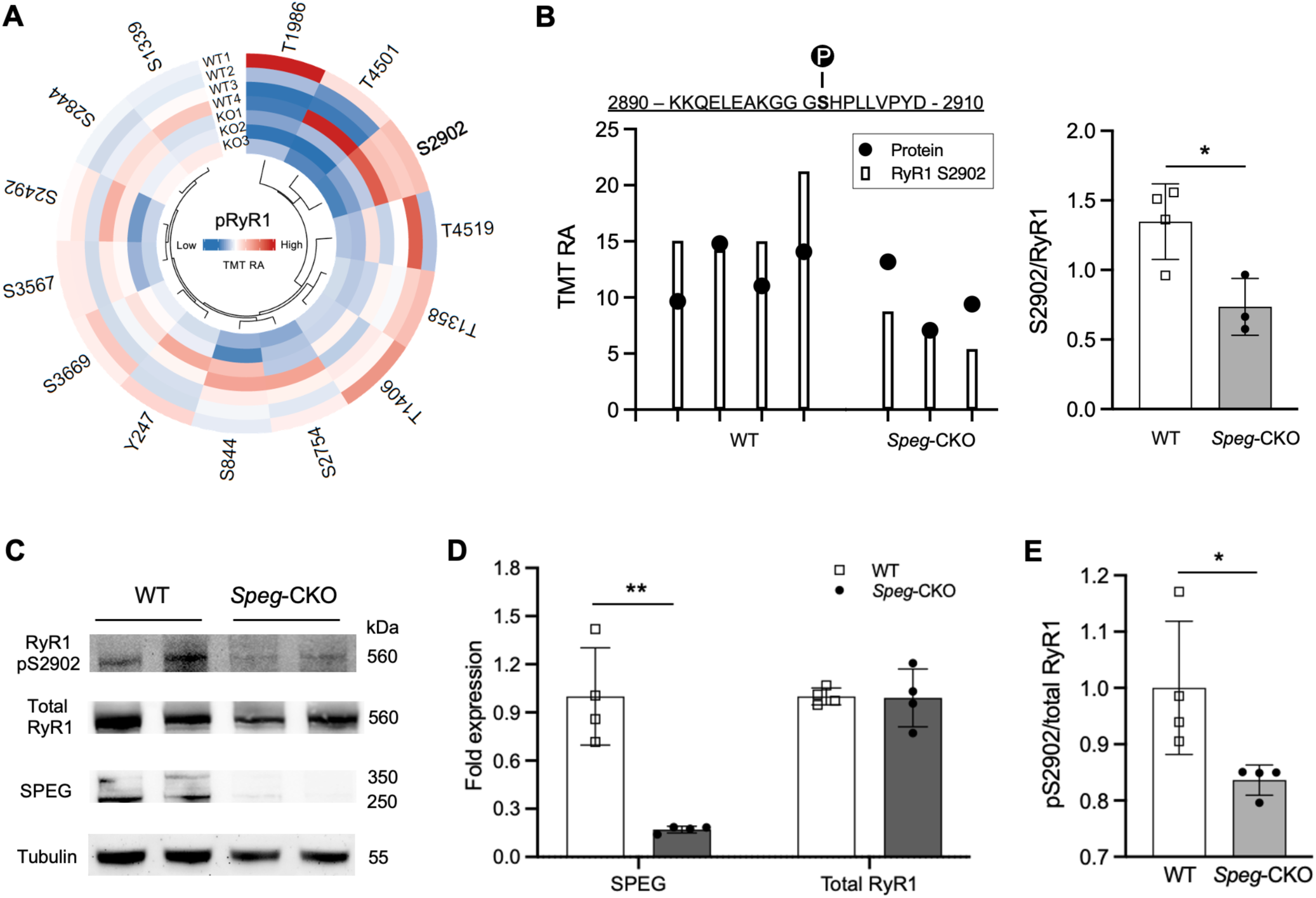
SPEG phosphorylates S2902 on RyR1 in skeletal muscle. (A) Circular heat map displaying the detected phosphosites of RyR1 by phosphoproteome. (B) RyR1 phosphoserine at 2902 (bars) and associated RyR1 protein levels (circles). Quantification of RyR1 S2902 levels normalized to total RyR1 protein level (*P < 0.05; unpaired 2-tailed t test). (C) Immunoblot image of SPEG, RyR1 S2902, total RyR1 protein levels in the skeletal muscle of WT and *Speg*-CKO mice. (D) Quantification of SPEG and RyR1 protein levels on immunoblot relative to tubulin (**P < 0.01, n = 4 per group; unpaired 2-tailed t test). (E) Quantification of RyR1 S2902 phosphorylation level on immunoblot relative to total RyR1 protein level (*P < 0.05, n = 4 per group; unpaired 2-tailed t test).

### Transcriptomic analysis indicates dysregulated pathways of ECM-receptor interaction and peroxisome proliferator-activated receptors (PPAR) signaling

To determine molecular pathways that are dysregulated in *Speg*-CKO mice, the quadriceps muscle transcriptomes of adult *Speg*-CKO (n =4) and WT mice (n =4) were assessed. PCA plots revealed that *Speg*-CKO groups separate from WT groups (**Figure 6A**), and volcano plots (**Figure 6B**) illustrated the differentially expressed genes (DEGs). Comparing *Speg*-CKO to WT mice, 312 DEGs were downregulated and 273 were upregulated (**Figure 6C**). The down- and upregulated DEGs were separately enriched by GO analysis (**Figure 6D**), and top 3 GO enrichments of BP, CC, and MF were listed in **Fig 6E**. ECM and fatty acid metabolism were the most affected GO in the transcriptome of *Speg*-CKO groups. Additionally, DEGs were also used for KEGG pathway analysis (**Fig 6F**), and pathways of ECM-receptor interaction and PPAR signaling were the most dysregulated in *Speg*-CKO groups. While genes linked to the ECM-receptor interaction pathway were generally downregulated (**Fig 6G and Supplemental Figure 2**), PPAR signaling pathway genes were predominantly upregulated (**Fig 6H**). Taken together, these findings suggest that SPEG deficiency affects the transcription of ECM and PPAR related genes in skeletal muscles.

**Figure 6.**
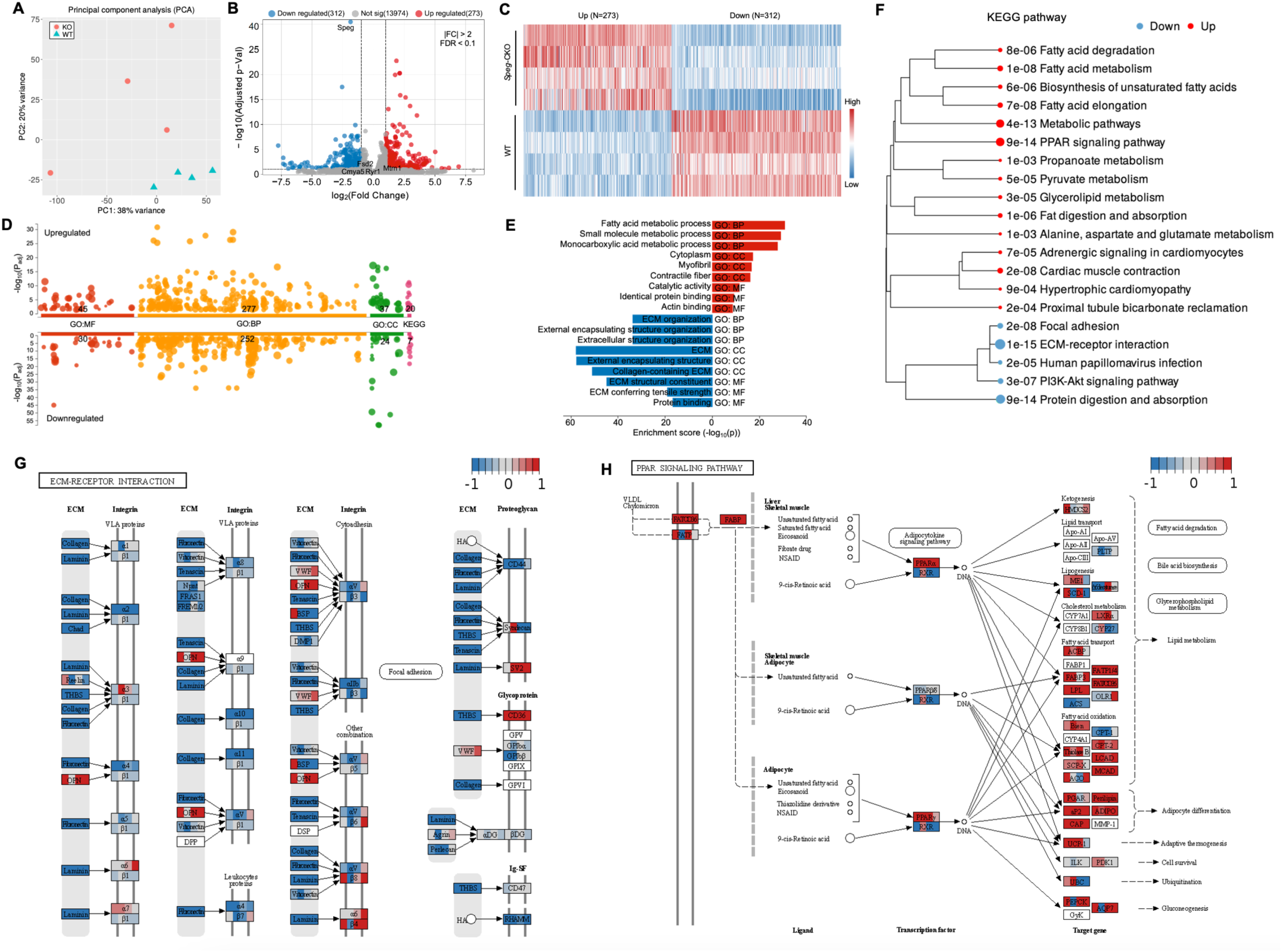
Transcriptome profiling in the skeletal muscle of *Speg*-CKO mice. (A) Principal component analysis of transcriptome data. The first and second axes are represented. Colored symbols represent genotypes for each mouse. (B) Volcano plots representing the differentially expressed genes (DEGs). Upregulated genes are in red, and downregulated genes are in blue (FDR <0.1 and a fold change beyond ±2). (C) Heat map of the genes that were differentially expressed in the skeletal muscle of *Speg*-CKO mice. (D) Enrichment analyses of upregulate and downregulated DEGs for GO and KEGG pathways. The number in the x-axis labels shows how many significantly enriched terms were found. The circle corresponds to term size. (E) Top three GO enrichment of BP, CC, and MF in (D). (F) Dendrogram displaying the KEGG enrichment. KEGG Pathview graphs of top downregulated (G) and upregulate DEGs (H). Upregulated genes are in red, and downregulated genes are in blue.

### Poor correlation between transcriptomic and proteomic data

The Pearson correlation analysis of transcriptomic and proteomic data did not reveal a strong correlation (0.22; **Supplemental Figure 3**) between detected transcripts and proteins, likely because of low expression and inconsistency between RNA and protein activity due to post-translational and other modifications (23, 24). For example, the transcriptomic data suggested that SPEG deficiency did not affect levels of myospryn complex (CMYA5, FSD2 and RyR1) in contrast to their considerable reduction of protein levels. This was further confirmed by the qPCR (**Supplemental Figure 4**) and immunoblot analysis.

## Discussion

We applied multi-omics approaches and follow up experiments to elucidate the fundamental role of SPEG in skeletal muscle function, critical for determining the molecular pathogenesis of SPEG-related disease and finding novel therapies. We have identified that SPEG is a novel binding partner of myospryn complex that include CMYA5, FSD2 and RyR1.

Recent studies have revealed that CMYA5 and FSD2 are concentrated at dyad in cardiac muscle and triad in skeletal muscle respectively, and they co-localize with ryanodine receptors at the junctional-SR (16). However, their role in skeletal muscle is poorly understood. CMYA5 is a large tripartite motif-containing protein (∼450 kD) highly expressed in skeletal and cardiac muscle, and it co-localizes with Z-lines, junctional sarcoplasmic reticulum proteins, and transverse tubules in mature cardiomyocytes (25). FSD2 (fibronectin type III and SPRY domain containing 2) is a protein (∼85 kD) closely related to the C-terminus of CMYA5. CMYA5 was previously reported to interact with various muscle proteins, including α-actinin, desmin, dysbindin, the RIIα regulatory subunit of PKA, dystrophin, calcineurin, titin, and calpain-3, with all these interactors binding to overlapping regions in the C-terminal tripartite motif of CMYA5 (16). Lu et al. reported that CMYA5 adjacent to Z-lines precedes junctional sarcoplasmic reticulum positioning during cardiac development, and its deficiency disrupts dyad architecture and junctional SR Ca^2+^ release, leading to cardiac dysfunction and inability to tolerate pressure overload (25). Interestingly, we have previously shown that SPEG deficiency in skeletal muscle leads to disruption of triad structure and calcium homeostasis with associated muscle dysfunction. In this study we have shown that SPEG interacts with myospryn complex, and its deficiency is associated with a significant reduction in amounts of myospryn complex proteins with unchanged transcript levels, indicating that SPEG plays a critical role in the regulation of myospryn complex in skeletal muscle.

Prior research has elucidated the role of SPEG in regulating calcium homeostasis in striated muscles. In cardiac muscle, SPEG can phosphorylate both RyR2^S2367^ (26) and SERCA2a^T484^ (27), which could inhibit diastolic calcium release and promote calcium reuptake into the SR. In skeletal muscle, we have previously shown that loss of SPEG affected the amplitude and the kinetics of RyR-mediated SR calcium release (9). Here, we further identified that RyR1-S2902 as a novel kinase substrate of SPEG and that SPEG loss alters the phosphorylation level of JPH2 at multiple sites. We hypothesize that loss of RyR1 phosphorylation due to SPEG deficiency decreases RyR1-mediated SR calcium release in the skeletal muscle.

We have previously demonstrated that SPEGβ interacts with desmin and DNM2 (14, 15), whereas SPEGα interacts with MTM1 using co-immunoprecipitation (6). Here, we found that antibody against FSD2 pulls down both SPEGα and SPEGβ, while RyR1 IP only pulls down SPEGα. There is no effective IP antibody for CMYA5, and further study is required to confirm their interaction. These findings indicate that SPEGα and SPEGb have unique targets suggestive of differential functions (**Figure 7**). It is interesting that patients with mutations affecting both SPEG isoforms are associated with a more severe clinical phenotype, while those with mutations affecting only SPEGβ are associated with a milder phenotype and often without cardiac involvement (7, 28). Indeed, SPEGα is essential for both skeletal and cardiac function and may partially compensate for SPEGβ deficiency. Overall, SPEGα and SPEGβ have unique interacting partners as summarized in **Table 1**, and we suspect that SPEGα and SPEGb have unique functions based on their interacting partners and patient phenotypes.

**Figure 7.**
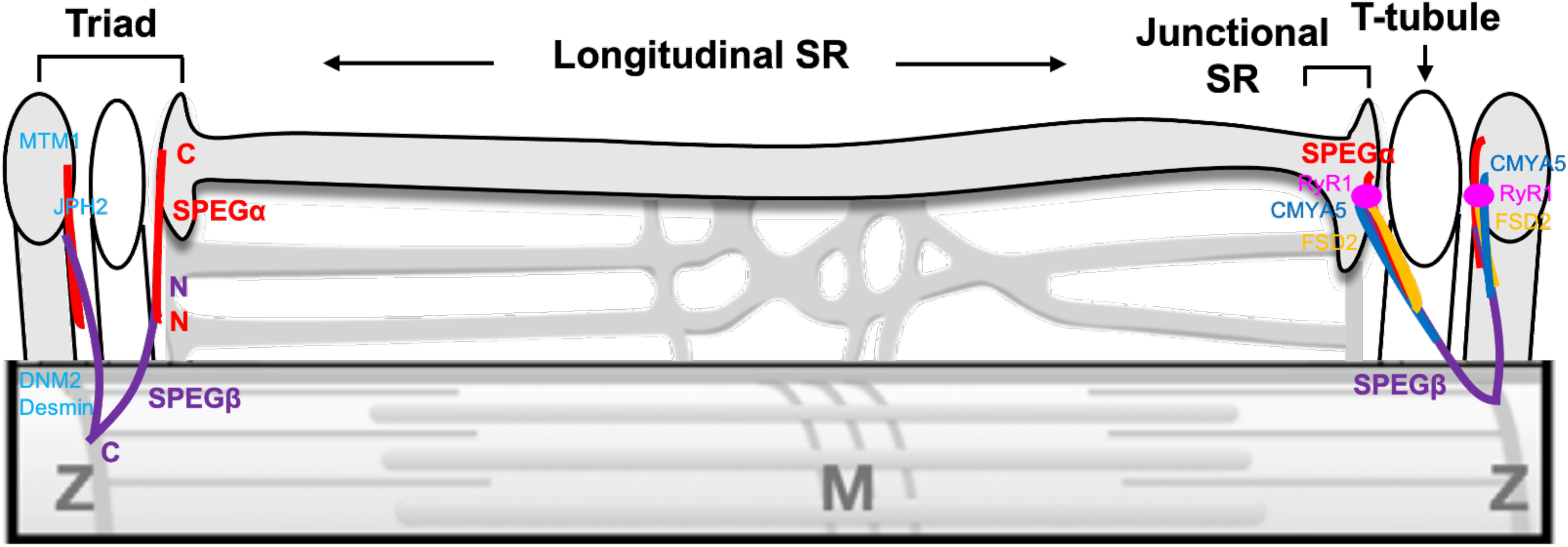
Hypothesis model of SPEG’s structural functions in skeletal muscles based on Table 1. Adapted from “Obscurin determines the architecture of the longitudinal sarcoplasmic reticulum” by Stephan Lange, 2009 (1). Red: SPEGα; purple: SPEGβ; C: C-terminus; N: N-terminus.

**Table 1.** SPEG and its interacting proteins in skeletal muscle.

| SPEG isoform | Region of SPEG | Interacting proteins | Protein location | Function in triad/dyad | Reference |
| --- | --- | --- | --- | --- | --- |
| SPEGα | Unknown | RyR1 | SR terminal cisternae | Calcium channel that mediates the release of Ca<sup>2+</sup> from the sarcoplasmic reticulum into the cytosol | This study |
|  | Ig like 9 and part of FnIII 2 domain (aa: 2530–2674) | MTM1<br>Phosphatase and coiled-coil domains (aa: 155-603) | SR terminal cisternae | T-tubule biogenesis and triad formation | [2] |
| SPEGβ | Ig like 9 and FnIII 2 domain (aa: 2200–2960) | Desmin<br>Rod domain (aa: 179-228) | Z line | Maintaining sarcomere structure, interconnecting Z-disks, and forming myofibrils, linking them to the sarcolemmal cytoskeleton, nucleus, and mitochondria | [3] |
|  | Unknown | DNM2 | Z line | T-tubule formation | [4] |
| Both isoforms | Unknown | CMYA5* | Both Z line and SR terminal cisternae | Assembly of ryanodine receptor clusters in striated muscle | This study |
|  | Unknown | FSD2 | Both Z line and SR terminal cisternae | Work with CMYA5 as myospryn complex | This study |

SR: sarcoplasmic reticulum; jSR: junctional sarcoplasmic reticulum; aa: amino acid; \* additional co-IP study will be required to confirm their interaction.

The mitochondrial abnormalities seen in SPEG-deficient muscles may be secondary to defects in triad and calcium homeostasis. In muscle cells, calcium is released from the SR, and defects in the triad can lead to abnormal calcium signaling that affect mitochondrial function. Calcium regulates key enzymes involved in mitochondrial oxidative phosphorylation, the process by which mitochondria generate ATP, and can also stimulate mitochondrial biogenesis and mitochondrial fusion (29). Recent findings show that interactions among SPEG, MTM1, DNM2, and BIN1 are essential for triad development and function (14, 30, 31). Triad and mitochondrial abnormalities have been seen with *Mtm1*-KO, *Bin1*-KO, and *Dnm2*-KI mice (**Supplemental Table 1**), and similarly, defective or absent triads and abnormally accumulated swollen mitochondria were detected in *Speg*-CKO muscle (32). Here, GSEA analysis of DEPs reveals upregulation of mitochondrial oxidoreductase, membrane protein complex, and lipid metabolism in *Speg*-CKO mice. Meanwhile, GO analysis of DEGs indicates significantly upregulated PPAR signaling pathway, fatty acid elongation and metabolism. PPARs are a class of nuclear receptors that play crucial roles in development and energy metabolism, and that they are master regulators of glucose and lipid homeostasis as well as modulators of mitochondrial function (33–35). Overall, the upregulation of PPAR signaling pathway and mitochondrial functions likely act to compensate for the triadic defects and calcium mishandling in the skeletal muscle of *Speg*-CKO mice.

In summary, we show that SPEG interacts with myospryn complex proteins in the skeletal muscle, and its deficiency results in myospryn complex abnormalities and decreased RyR1 phosphorylation level at S2902. Additionally, we identify that SPEG phosphorylates multiple phosphorylation sites of JPH2 suggests critical phosphorylation function of kinase domains of SPEG. We also determined that SPEGα and SPEGβ have unique interacting partners which suggests their differential function needing further elucidation. Further, the mitochondrial defects seen in *Speg*-CKO skeletal muscle may be a consequence of abnormal calcium signaling. This study demonstrates the advantages of using multi-omics techniques to comprehensively analyze the pathologic and molecular anomalies of rare diseases, setting the groundwork for the development of novel and precise therapeutics for patients carrying *SPEG* mutations.

## Materials and Methods

### Animal model

All studies were approved by the Institutional Animal Care and Use Committee at Children’s Hospital Boston (approval number 20-05-4179). The work followed the Guide for the Care and Use of Laboratory Animals and all the regulatory protocols set forth by the Boston Children’s Hospital Animal Resources at Children’s Hospital (ARCH) facility. *Speg*-CKO mice have been generated as previously described (36). Homozygous *Speg*-conditional KO mice (*Speg^fl/fl^*) were bred with male transgenic mice who have the Cre recombinase driven by muscle creatine kinase promoter (MCK-Cre^+^), with Cre activity observed in skeletal and cardiac muscle. Specific primers were used to identify *Speg^fl/fl^* and MCK-Cre^+^ alleles (36, 37). In the figure legends, each experiment’s sample size is specified.

### Quantitative PCR with reverse transcription (RT–qPCR)

Total RNA was isolated from skeletal muscles of WT and *Speg*-CKO mice using the mirVana miRNA Isolation Kit (AM1561, Thermofisher) according to manufacturer’s protocol. One µg of total RNA was used to generate complementary DNA (cDNA) using the SuperScript™ IV First-Strand Synthesis System and random hexamers (cat# 18091050, ThermoFisher Scientific). qRT-PCR was performed using PowerUp SYBR Green Master mix (A25742, Thermofisher) in a QuantStudio™ 3 Real-Time PCR System. Real-time quantitative PCR (qRT-PCR) primers were listed in **Supplemental Table 2**.

### Immunoblot analysis

Skeletal muscles from WT, *Speg*-CKO, and *Mtm1*-KO mice were dissected, snap frozen in isopentane, and stored at −80°C until analysis. Protein isolation and western blot procedures were performed as described previously (38). Immunofluorescent western blot was performed. Proteins were probed with primary antibodies against rabbit anti-SPEG (12472-T16, 1:1000 dilution, SinoBiological, Beijing, China), mouse anti-RyR1 (sc-376507, 1:800 dilution, Santa Cruz Biotechnology), rabbit anti-CMYA5 (1:800 dilution, from Dr. Francisco J. Naya, Boston University), rabbit anti-MTM1 (PI168, 1:800 dilution, from IGBMC), mouse anti-FSD2 (sc-393072, 1:1000 dilution, Santa Cruz Biotechnology), and custom rabbit-anti-RyR1-pS2902 (YenZym, South San Francisco, CA; 1:800). IRDye 800CW Donkey anti-Rabbit IgG Secondary antibody (926-32213, 1:5000, LI-COR), IRDye 680RD Donkey anti-Mouse IgG Secondary antibody (926-68072, 1:5000, LI-COR), and anti-tubulin Rhodamine antibody (AbD22584, 1:5000, Bio-Rad Laboratories, Hercules, CA, USA) were used for immunofluorescence detection. Quantification of protein levels normalized to tubulin was performed using ImageJ software.

### Co-immunoprecipitation (co-IP)

Skeletal muscle lysates from quadriceps, gastrocnemius, and triceps were obtained by homogenization via Cryogrinder and lysed in Pierce IP lysis buffer (PI87787, Thermofisher) supplemented with Halt Protease and Phosphatase Inhibitor (PI78441, Thermofisher) and 5mM EDTA (final concentration). Tissue was lysed at 4°C for 30 minutes. After centrifugation (16,000*g*, 4°C, 20 minutes), the soluble fractions were collected, and the concentration was measured using a colorimetric BCA assay (23225; Thermo Fisher Scientific). Soluble homogenates were precleared with Dynabead Protein G beads (Thermo Fisher Scientific) for 1 hour, and supernatants were incubated with specific antibodies directed against the protein of interest at 4°C for 12 to 24 hours. Dynabead Protein G beads were then added for 2 hours to capture the immune complex. Beads were washed 3 times with co-IP buffer supplemented with 0.1% CHAPS. For all experiments, 2 negative controls consisted of a sample lacking the primary antibody and a sample incubated with another primary antibody from the same serotype as the antibody of interest. Resulting beads were eluted with Laemmli buffer and subjected to SDS-PAGE followed by immunoblot.

### Identification of the interactome

Immunoprecipitation of SPEG was performed on pooled skeletal muscles from at least 3 WT mice for each experiment. A rabbit anti-SPEG antibody (12472-T16, 1:50 dilution, SinoBiological, Beijing, China) was used for immunoprecipitation. Samples were concentrated and analyzed by SDS–PAGE and stained with the Coomassie blue (1610786, Bio-Rad Laboratories, Hercules, CA, USA). The IgG control and the *Speg*-CKO lysate incubated with SPEG antibody were used as controls. Excised gel bands were cut into approximately 1 mm^3^ pieces. Gel pieces were then subjected to a modified in-gel trypsin digestion procedure (39). Gel pieces were washed and dehydrated with acetonitrile for 10 min. followed by removal of acetonitrile. Pieces were then completely dried in a speed-vac. Rehydration of the gel pieces was with 50 mM ammonium bicarbonate solution containing 12.5 ng/µl modified sequencing-grade trypsin (Promega, Madison, WI) at 4°C. After 45 min., the excess trypsin solution was removed and replaced with 50 mM ammonium bicarbonate solution to just cover the gel pieces. Samples were then placed in a 37°C room overnight. Peptides were later extracted by removing the ammonium bicarbonate solution, followed by one wash with a solution containing 50% acetonitrile and 1% formic acid. The extracts were then dried in a speed-vac (∼1 hour).

The samples were reconstituted in 5 - 10 µl of HPLC solvent A (2.5% acetonitrile, 0.1% formic acid). A nano-scale reverse-phase HPLC capillary column was created by packing 2.6 µm C18 spherical silica beads into a fused silica capillary (100 µm inner diameter x ∼30 cm length) with a flame-drawn tip (40). After equilibrating the column each sample was loaded via a Famos auto sampler (LC Packings, San Francisco CA) onto the column. A gradient was formed, and peptides were eluted with increasing concentrations of solvent B (97.5% acetonitrile, 0.1% formic acid). As peptides eluted, they were subjected to electrospray ionization and then entered an LTQ Orbitrap Velos Pro ion-trap mass spectrometer (Thermo Fisher Scientific, Waltham, MA). Peptides were detected, isolated, and fragmented to produce a tandem mass spectrum of specific fragment ions for each peptide. Peptide sequences (and hence protein identity) were determined by matching protein databases with the acquired fragmentation pattern by the software program, Sequest (Thermo Fisher Scientific, Waltham, MA) (41). All databases include a reversed version of all the sequences and the data was filtered to between a one and two percent peptide false discovery rate.

### Transmission electron microscopy (EM)

Skeletal muscle samples of WT and *Speg*-CKO quadriceps (1–2 mm cubes, n = 3 per group) were fixed in 2.5% glutaraldehyde, 1.25% paraformaldehyde and 0.03% picric acid in 0.1 M sodium cacodylate buffer (pH 7.4) overnight in room temperature and stored at 4 °C. They were then washed in 0.1 M cacodylate buffer and post-fixed with 1% osmium tetroxide (OsO_4_)/1.5% potassium ferrocyanide (KFeCN6) for 1h, washed in water 3× and incubated in 1% aqueous uranyl acetate for 1 h followed by two washes in water and subsequent dehydration in grades of alcohol (10 min each; 50, 70, 90%, 2 × 10 min 100%). The samples were then put in propylene oxide for 1h and infiltrated overnight in a 1:1 mixture of propylene oxide and TAAB Epon (Marivac Canada Inc., St Laurent, Canada). The following day, the samples were embedded in TAAB Epon and polymerized at 60°C for 48h. Ultrathin sections (∼60 nm) were cut on a Reichert Ultracut-S microtome, picked up on to copper grids stained with lead citrate and examined in a JEOL 1200EX Transmission electron microscope, and images were recorded with an AMT 2k CCD camera. This was performed at the EM Core of the Harvard Medical School.

### Phosphoproteome profiling

Skeletal muscle samples of WT (n = 4) and *Speg*-CKO (n = 3) mice were lysed and processed as described in the SPEED protocol (42). Following digestion with Lys-C and trypsin, peptides were desalted by using 100mg SepPak columns. The elutions were dried via vacuum centrifugation and the phosphopeptides were enriched with the High-Select Fe^3+^-NTA Phosphopeptide Enrichment Kit according to manufacturer’s specifications using approximately 1 mg protein digest per enrichment column. The elutions were dried via vacuum centrifugation, while the flow-throughs were saved for subsequent whole proteome analysis.

TMT labeling. In general, we estimate that at most 10 ug of phosphopeptides were enriched from 1 mg of total peptide. We added 20 ug of TMTpro reagent (Thermo-Fisher; Lot #:WL338745) to the peptides (∼10 µg) along with acetonitrile to again achieve a final acetonitrile concentration of approximately 30% (v/v) in a total volume of 50 µL. Following incubation at room temperature for 1 h, the reaction was quenched with hydroxylamine to a final concentration of 0.3% (v/v). The sample was vacuum centrifuged to near dryness and subjected to C18 solid-phase extraction (SPE, Sep-Pak).

Off-line basic pH reversed-phase (BPRP) fractionation. We fractionated the peptide samples using BPRP HPLC. We used an Agilent 1200 pump equipped with a degasser and a UV detector. Peptides were subjected to a 50-min linear gradient from 5% to 35% acetonitrile in 10 mM ammonium bicarbonate pH 8 at a flow rate of 0.6 mL/min over an Agilent 300Extend C18 column (3.5 μm particles, 4.6 mm ID and 250 mm in length). The peptide mixture was fractionated into a total of 96 fractions, which were consolidated into 24 super-fractions (in a checkerboard-like pattern). Samples were subsequently acidified with 1% formic acid and vacuum centrifuged to near dryness. Each consolidated fraction was desalted by StageTip, and reconstituted in 5% acetonitrile, 5% formic acid for LC-MS/MS processing.

Mass spectrometric data were collected on an Orbitrap Eclipse mass spectrometer coupled to a Proxeon NanoLC-1200 UHPLC (ThermoFisher Scientific). The 100 µm capillary column was packed in-house with 35 cm of Accucore 150 resin (2.6 μm, 150Å; ThermoFisher Scientific). Data were acquired for 120 min per fraction. The scan sequence began with an MS1 spectrum: Orbitrap analysis, resolution 120,000, 400−1500 Th, automatic gain control (AGC) target 400K, maximum injection time 50 ms. MS2 analysis, which occurred in the OrbiTrap, consisted of higher-energy collision dissociation (HCD), AGC 150K, NCE (normalized collision energy) 36, isolation window 0.5 Th, maximum injection time set to 250 and TopSpeed set at 1.5 sec. Each of the samples were shot twice using two sets of compensation voltages. For FAIMS, the dispersion voltage (DV) was held constant at 5000V, the compensation voltages (CVs) were set at −35V, −55V, and −75V for the first shot and −45 and −65V for the second shot. In total each of the 12 fractions were run twice, for a total of 24 runs.

Database searching included all entries from the mouse UniProt Database (downloaded: 2021). The database was concatenated with one composed of all protein sequences for that database in the reversed order (43). Raw files were converted to mzXML, and monoisotopic peaks were re-assigned using Monocle (44). Searches were performed with Comet (45) using a 50-ppm precursor ion tolerance for total protein level profiling. The product ion tolerance was set to 0.02 Da. TMTpro labels on lysine residues and peptide N-termini (+304.207 Da), as well as carbamidomethylation of cysteine residues (+57.021 Da) were set as static modifications, while oxidation of methionine residues (+15.995 Da), phosphorylation (+79.966), and deamidation (+0.984) were set as variable modifications. Peptide-spectrum matches (PSMs) were adjusted to a 1% false discovery rate (FDR) using a linear discriminant after which proteins were assembled further to a final protein-level FDR of 1% analysis (46). AScore was used to determine site localization (47), with a score of 13 denoting 95% confidence for a specified phosphorylation site. Phosphorylation sites were quantified by summing reporter ion counts across all matching PSMs. More specifically, reporter ion intensities were adjusted to correct for the isotopic impurities of the different TMTpro reagents according to manufacturer specifications. Peptides were filtered to include only those with a summed signal-to-noise (SN) ≥ 100 across all TMT channels. An extra filter of an isolation specificity (“isolation purity”) of at least 0.5 in the MS1 isolation window was applied for the phosphorylated peptide analysis. The signal-to-noise (S/N) measurements of peptides were globally normalized using the protein normalization factors mentioned below to account for equal protein loading. Cutoff values for differentially expressed phosphorylation peptide determinations were as follows: p value <0.05 and absolute value of log2FC >1.5.

### Whole proteome profiling

Flow-throughs from the phospho-enrichment described above, 50 µg per replicate, were used for the whole proteome work for each sample. 120 µg of TMTpro reagents (Thermo-Fisher; Lot #:WL338745) were added to the peptides (50 µg) along with acetonitrile to achieve a final acetonitrile concentration of approximately 30% (v/v) in a total volume of 100 µL. Following incubation at room temperature for 1 h, the reaction was quenched with hydroxylamine to a final concentration of 0.3% (v/v). The sample was vacuum centrifuged to near dryness and subjected to C18 solid-phase extraction (SPE, Sep-Pak). Off-line basic pH reversed-phase (BPRP) fractionation was the same as phosphoproteome profiling.

Mass spectrometric data were collected on an Orbitrap Fusion Lumos mass spectrometer coupled to a Proxeon NanoLC-1200 UHPLC (ThermoFisher Scientific). A 100 µm capillary column was packed in-house with 35 cm of Accucore 150 resin (2.6 μm, 150Å; ThermoFisher Scientific). Data were acquired for 90 min per fraction. The scan sequence began with an MS1 spectrum: Orbitrap analysis, resolution 60,000, 400−1600 Th, automatic gain control (AGC) target 400K, maximum injection time 50 ms. MS2 analysis consisted of collision-induced dissociation (CID), quadrupole ion trap analysis, AGC 10K, NCE (normalized collision energy) 35, q-value 0.25, isolation window 0.6 Th, maximum injection time set to 35 and TopSpeed set at 1.25 sec. An on-line real-time search algorithm (Orbiter) was used to trigger MS3 scans for quantification (48). For the MS3 scan (performed in the OrbiTrap), we used higher-energy collision dissociation (HCD) with NCE 55%, AGC 200K, maximum injection time 200 ms, resolution 50,000 at 400 Th, isolation window 1.2. The close out was set at two peptides per protein per fraction (48). For High-field Asymmetric-waveform Ion Mobility spectrometry (FAIMS), the dispersion voltage (DV) was held constant at 5000V, the compensation voltages (CVs) were set at −40V, −60V, and −80V (49). In total 24 fractions were analyzed for each multiplexed experiment.

Searches were performed as described above but with product ion tolerance was set to 0.9 Da (as the MS2 scans are low-resolution) and without phosphorylation or deamidation variable modifications. Proteins were quantified by summing reporter ion counts across all matching PSMs. More specifically, reporter ion intensities were adjusted to correct for the isotopic impurities of the different TMTpro reagents according to manufacturer specifications. Peptides were filtered to include only those with a summed signal-to-noise (SN) ≥ 100 across all TMT channels. The signal-to-noise (S/N) measurements of peptides assigned to each protein were summed (for a given protein). These values were normalized so that the sum of the signal for all proteins in each channel was equivalent thereby accounting for equal protein loading. The resulting normalization factors will be used to normalize the phosphorylation sites as discussed above to account for equal protein loading. Finally, each protein abundance measurement was scaled, such that the summed signal-to-noise for that protein across all channels equals 100, thereby generating a relative abundance (RA) measurement. Cutoff values for differentially expressed protein (DEPs) determinations were as follows: p value <0.05 and absolute value of fold change >1.5.

### Transcriptome profiling

Skeletal muscles from quadriceps femoris were isolated from WT and *Speg*-CKO mice. A total of 8 mRNA samples (n = 4 for per group) was isolated and was then converted into double-stranded DNA (dsDNA). Procedures including the sample quality control (QC), the library preparation, and sequencing for all the samples were performed by Novogene (Sacramento, CA, USA). RNA-seq raw data were aligned using “align” (Rsubread, V2.0.1) and the latest UCSC mouse annotation (GRCm38/mm10). The raw data were trimmed before aligned using Trimmomatic (V0.39) for QC. Gene-level read counts were quantified using “featureCounts” (Rsubread, V2.0.1). To identify differentially expressed genes, DESeq2 (version 1.26.0) was used with default parameters in Bioconductor packages. All software/packages were run using their default parameters. Count tables were normalized to TPM (Transcripts per Million) for visualizations and QC. Sample clustering and standard path analyses (GO and KEGG) were performed using a custom-made pipeline (VExP-RNseq). Transcripts were called as differentially expressed when the adjusted p values were below 0.1 and fold-changes were over ±2.

### Bioinformatic analysis of omics data

For proteome and transcriptome analysis, DEPs and DEGs both were analyzed for GO and KEGG using g:Profiler (version e107_eg54_p17_bf42210) (50). Additionally, proteome data was conducted the unbiased Gene Set Enrichment Analysis (GSEA), and all GSEA plots including GSEA enrichment plot and enrichment map were generated with the GSEA software (version 4.2.1) (51). The enrichment map was exported with CYTOSCAPE (version 3.9.1) (52). Plots of KEGG pathways were generated using KEGG Mapper (version 5.0) (53).

### Statistical analysis

Results were analyzed with GraphPad Prism (v.8.0; GraphPad Software) and expressed as mean ± standard deviation (SD). Unpaired 2-tailed *t* test was used to determine statistically significant differences for 2-group comparisons. One-way ANOVA followed by Tukey’s post hoc test was used for multiple-group comparisons. The numbers of samples per group (*n*) and statistical significance for all comparisons are specified in the figure legends. *P* < 0.05 was considered statistically significant.

## Data availability

The mass spectrometry data (interactome, proteome, and phosphoproteome) have been deposited to the ProteomeXchange Consortium via the PRIDE partner repository with the dataset identifier PXD041692. RNA-seq data were deposited in NCBI SRA: SUB12947820.

## Acknowledgements

PBA was supported by R01 AR068429 from the National Institute of Arthritis and Musculoskeletal and Skin Diseases of National Institute of Health (NIH). This work was supported in part by the resources of the IDDRC Molecular Genetics Core funded by U54HD090255 from the National Institutes of Health. The authors would like to thank Dr. Francisco J. Naya at Boston University and Dr. William Pu and Dr. Fujian Lu at Boston Children’s Hospital for providing CMYA5 antibody as well as Dr. Jocelyn Laporte at Institut de Génétique et de Biologie Moléculaire et Cellulaire (IGBMC) for providing MTM1 antibody.

Mass spectrometry-based protein–protein interaction was performed in the Harvard Medical School Taplin Mass Spectrometry Facility. Proteome and phosphoproteome experiments were conducted in the Harvard Medical School Thermo Fisher Scientific Center for Multiplexed Proteomics. Electron microscopy imaging, consultation, and/or services were performed in the Harvard Medical School Electron Microscopy Facility. The graphical abstract was created with BioRender.com. The funder had no role in study design, data collection and analysis, decision to publish, or preparation of the manuscript.

## Author contributions

QL and PBA designed the experiments and project administration. QL, JL, SL, KSA, RA, MM, BM, AHB, XL, MAP, and PBA carried out experiments, performed data analyses, and drafted the manuscript. All authors read and approved the final manuscript.

## Conflict of interest

The authors declare that they have no conflict of interest.

## Notes

### Competing Interest Statement

The authors have declared no competing interest.

